# Metagenomic analysis unravels novel taxonomic differences in the uterine microbiome between healthy mares and mares with endometritis

**DOI:** 10.1101/2022.10.09.511479

**Authors:** Aeknath Virendra, Sarita U. Gulavane, Zulfikar A. Ahmed, Ravi Reddy, Ravindra J. Chaudhari, Sandeep M. Gaikwad, Raju R. Shelar, Shailesh D. Ingole, Varsha D. Thorat, Afroza Khanam, Firdous Ahmad Khan

## Abstract

The application of high throughput technologies has enabled unravelling of unique differences between healthy mares and mares with endometritis at transcriptomic and proteomic levels. However, differences in the uterine microbiome are yet to be investigated. The present study was aimed at evaluating the differences in uterine microbiome between healthy mares and mares with endometritis. Low-volume lavage (LVL) samples were collected from the uterus of 30 mares classified into healthy (n=15) and endometritis (n=15) based on their reproductive history, intrauterine fluid accumulation, gross appearance of LVL samples, endometrial cytology, and bacterial culture. The samples were subjected to metagenomic analysis using 16S rRNA sequencing. Notable differences in the uterine microbiome were observed between healthy mares and mares with endometritis at various taxonomic levels. In healthy mares, the most abundant phylum, class, order, and family were Firmicutes, Bacilli, Bacillales, and Paenibacillaceae, respectively. In contrast, the most abundant corresponding taxonomic levels in mares with endometritis were Proteobacteria, Gammaproteobacteria, Enterobacterales, and Enterobacteriaceae, respectively. At the genus level, *Brevibacillus* and *Paenibacillus* were more abundant in healthy mares, whereas *Escherichia, Salmonella*, and *Klebsiella* were more abundant in mares with endometritis. In healthy mares, *Brevibacillus brevis* was the most abundant species, followed by *Brevibacillus choshinensis* and *Paenibacillus sp JDR-2*. However, in mares with endometritis, *Escherichia coli* was the most abundant species, followed by *Salmonella enterica* and *Klebsiella pneumoniae*. These results indicate novel differences in the uterine microbiome between healthy mares and mares with endometritis. The findings can potentially help formulate new approaches to prevent or treat equine endometritis.

## 1. Introduction

Endometritis is one of the major problems in the equine industry. In a survey of equine practitioners, endometritis was ranked as the third most common medical problem in equine practice after colic and viral respiratory tract disease (Traub-Dargatz et al., 1991). About 25 to 60% of barren mares were shown to have endometritis in different studies (Collins, 1964; Bain, 1966; Morris and Allen, 2002). Equine breeders suffer severe economic losses annually due to reduced pregnancy rates associated with endometritis (Riddle et al., 2007) combined with the diagnostic, therapeutic and rebreeding costs (LeBlanc, 2010). Despite several decades of research, mechanisms underlying equine endometritis are yet to be fully elucidated. The application of high throughput technologies in the recent years has advanced our understanding of the differences between uterine health and disease at genomic (Weber et al., 2021) and proteomic (Diel de Amorim et al., 2020) levels. Similar studies on the uterine microbiome are required to elucidate the etiological basis of endometritis and the variations in predisposition to this condition between different mares.

The dogma that the healthy uterus is sterile has been challenged by research studies conducted in various species including cattle (Moore et al., 2017), horses (Heil et al., 2018; Holyoak et al., 2022), and dogs (Lyman et al., 2019). Several bovine studies have demonstrated that the uterine microbiome is dynamic with remarkable differences reported between a healthy state and disease conditions such as endometritis (Bicalho et al., 2017; Pascottini et al., 2020; Ballas et al., 2021). To our knowledge, differences in the uterine microbiome between health and disease in mares have not been investigated yet. Therefore, the objective of the present study was to evaluate the differences in uterine microbiome between healthy mares and mares with endometritis.

## 2. Material and Methods

This study was conducted at the Department of Animal Reproduction, Gynecology and Obstetrics, Mumbai Veterinary College located in Maharashtra, India. All animal handling and clinical procedures used in this study were approved by the Institutional Ethics Committee for Veterinary Clinical Research (BVC/IEC-VCR/2021/dated 15/06/2021).

### 2.1. Animals

A total of 48 Thoroughbred mares (5 to 12 years old, BCS 5 to 6) from three separate stud farms in and around Pune, India were examined to select 15 healthy mares and 15 mares with endometritis. The classification into healthy mares and mares with endometritis was based on breeding history and results of transrectal examination, low-volume lavage characteristics, endometrial cytology, and bacterial culture. Diagnosis of endometritis was based on a history of infertility, excessive intrauterine fluid accumulation (> 2 cm) noted on transrectal examinations, turbidity of low-volume uterine lavage samples, inflammatory endometrial cytology (greater than 2 neutrophils per high power field), and positive bacterial culture (LeBlanc and Causey, 2009; Katila, 2016).

### 2.2. Collection of low-volume uterine lavage samples

Sample collection was performed during the period from January to September, 2021.

Mares were sampled during estrus, detected on the basis of behavioral signs and results of transrectal examinations. After restraining each mare in stocks, the tail was wrapped, and the perineum was cleaned using soap and water. A low-volume uterine lavage sample was collected transcervically from each mare using 100 mL sterile water and a sterile lavage assembly according to the procedure reported previously (Maloney et al., 2019). The samples were stored at − 80°C until DNA extraction for metagenomic analysis. The samples from each group were pooled randomly using Microsoft Excel randomization function (pooled samples E1, E2, E3, and E4 from mares with endometritis and pooled samples H1, H2, and H3 from healthy mares).

### 2.3. Metagenomic analysis

#### 2.3.1. Library construction and sequencing

Total genomic DNA was extracted using the Qiagen DNeasy mini-spin column (DNeasy Mini Kit, Qiagen, USA) according to the manufacturer’s instructions. The extracted DNA was quantified using NanoDrop 1000 spectrophotometer (ThermoFisher Scientific, Waltham, MA, USA). For each PCR reaction, 20 ng of total DNA was used as a template. Amplification was performed with primers specific to V3-V4 regions of the bacterial 16S rRNA gene fragment (Forward 5’-CCTAYGGGRBGCASCAG-3’, Reverse 5’-GGACTACNNGGGTATCTAAT-3’). The amplification of 16S rRNA genes was conducted using the KAPA2G™ Robust HotStart ReadyMix PCR Kit (Kapa Biosystems, Wilmington, MA, USA) in a total volume of 25□μl. Amplification was performed with the following PCR conditions: initial denaturation at 95□°C for 3□min, 5□cycles of 95□°C for 15□s, 55□°C for 15 □s, and 72□°C for 30□ s, 30□cycles of 95□°C for 15□s, 62□°C for 15□s, and 72□°C for 30□s, followed by a final extension at 72□°C for 1□min using the grade PCR instrument BioRad T-100. Amplified DNA was purified using AMPure® XP (Beckman Coulter) and quantified by a NanoDrop® 1000 (ThermoFisher Scientific, Waltham, MA, USA). Illumina TruSeq DNA PCR-free Library Preparation Kits (Illumina, San Diego, CA, USA) were used for library sequencing according to the manufacturer’s protocol. Index codes were added to all the samples. The Qubit 2.0 Fluorometer (ThermoFisher Scientific, Waltham, MA, USA) and the Agilent Bioanalyzer systems were used to assess library quality and the 250 bp paired-end reads were obtained after sequencing on the Illumina HiSeq platform (Illumina, San Diego, CA, USA).

#### 2.3.2. Metagenome assembly, mapping, and taxonomical assignment

MEGAHIT (Version 1.1.3), an assembler that can assemble large and complex metagenomics data, was used to assemble the processed reads. Minimum multiplicity for filtering (k_min+1)-mers was set as 2 and Minimum length of contigs to output was 200. The assembled reads were used for mapping using Mega BLAST against target database (nt_2018-01-22). Kraken2 (Version 2.1.1) was applied for taxonomical classification using the Kraken2 database pluspf2021-05.

#### 2.3.3. Statistical analysis

Data was analyzed using QIIME (Quantitative Insights into Microbial Ecology).

Alpha diversity indices (Simpson, Shannon, and Chao1) were calculated to assess the variability of species within a sample while beta diversity was calculated using Bray-Curtis dissimilarity to assess the differences in composition between samples. Principal component analysis (PCA) plot was used to visualize the variance between the samples.

## 3. Results

Metagenomic analysis revealed the existence of diverse microbial communities in uterine samples from all mares included in the study (Tables 1 and 2). As shown in the PCA plot, there was a difference in the microbiome composition between healthy mares and mares with endometritis (Figure 1). The relative abundance of the various taxonomic levels in the two groups of mares is shown in Figure 2 (panels a-g). The most abundant kingdom in both groups was Bacteria (Figure 2a). In healthy mares, Firmicutes was the major phylum followed by Proteobacteria. There was an exactly opposite trend in mares with endometritis where the major phylum was Proteobacteria followed by Firmicutes (Figure 2b). Bacilli were found to be the most abundant class in healthy mares, whereas Gamma Proteobacteria were the most abundant class in mares with endometritis (Figure 2c). The most abundant order in healthy mares was Bacillales followed by Lactobacillales and Clostridiales, whereas in mares with endometritis, the most abundant order was Enterobacterales followed by Xanthomonadales and Pseudomonadales (Figure 2d). Paenibacillaceae was the most abundant family in healthy mares followed by Streptococcaceae and Clostridiaceae. On the other hand, in mares with endometritis, the most abundant family was Enterobacteriaceae followed by Xanthomonadaceae and Moraxellaceae (Figure 2e).

**Table 1.** Alpha diversity indices of the microbiome in uterine samples from healthy mares (H1-H3) and mares with endometritis (E1-E4)

| Sample | Simpson (1-D) | Shannon (H) | Chao1 |
| --- | --- | --- | --- |
| H1 | 0.87 | 2.51 | 1827 |
| H2 | 0.95 | 3.95 | 2388 |
| H3 | 0.95 | 3.85 | 2113 |
| E1 | 0.87 | 2.38 | 1886 |
| E2 | 0.89 | 2.65 | 2083 |
| E3 | 0.88 | 2.49 | 2042 |
| E4 | 0.88 | 2.58 | 1868 |

**Table 2.**
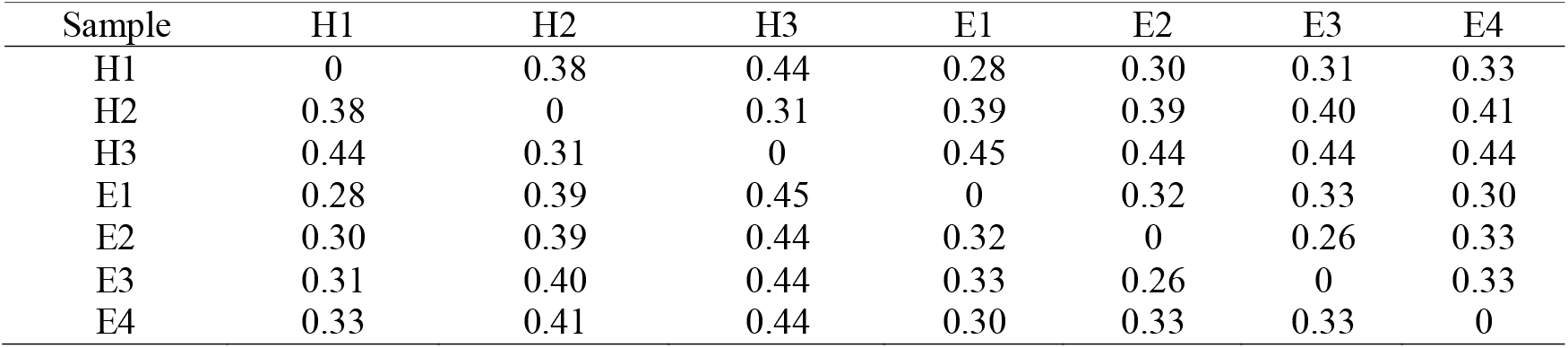
Beta diversity of the microbiome in uterine samples from healthy mares (H1-H3) and mares with endometritis (E1-E4)

| Sample | H1 | H2 | H3 | E1 | E2 | E3 | E4 |
| --- | --- | --- | --- | --- | --- | --- | --- |
| H1 | 0 | 0.38 | 0.44 | 0.28 | 0.30 | 0.31 | 0.33 |
| H2 | 0.38 | 0 | 0.31 | 0.39 | 0.39 | 0.40 | 0.41 |
| H3 | 0.44 | 0.31 | 0 | 0.45 | 0.44 | 0.44 | 0.44 |
| E1 | 0.28 | 0.39 | 0.45 | 0 | 0.32 | 0.33 | 0.30 |
| E2 | 0.30 | 0.39 | 0.44 | 0.32 | 0 | 0.26 | 0.33 |
| E3 | 0.31 | 0.40 | 0.44 | 0.33 | 0.26 | 0 | 0.33 |
| E4 | 0.33 | 0.41 | 0.44 | 0.30 | 0.33 | 0.33 | 0 |

**Figure 1.**
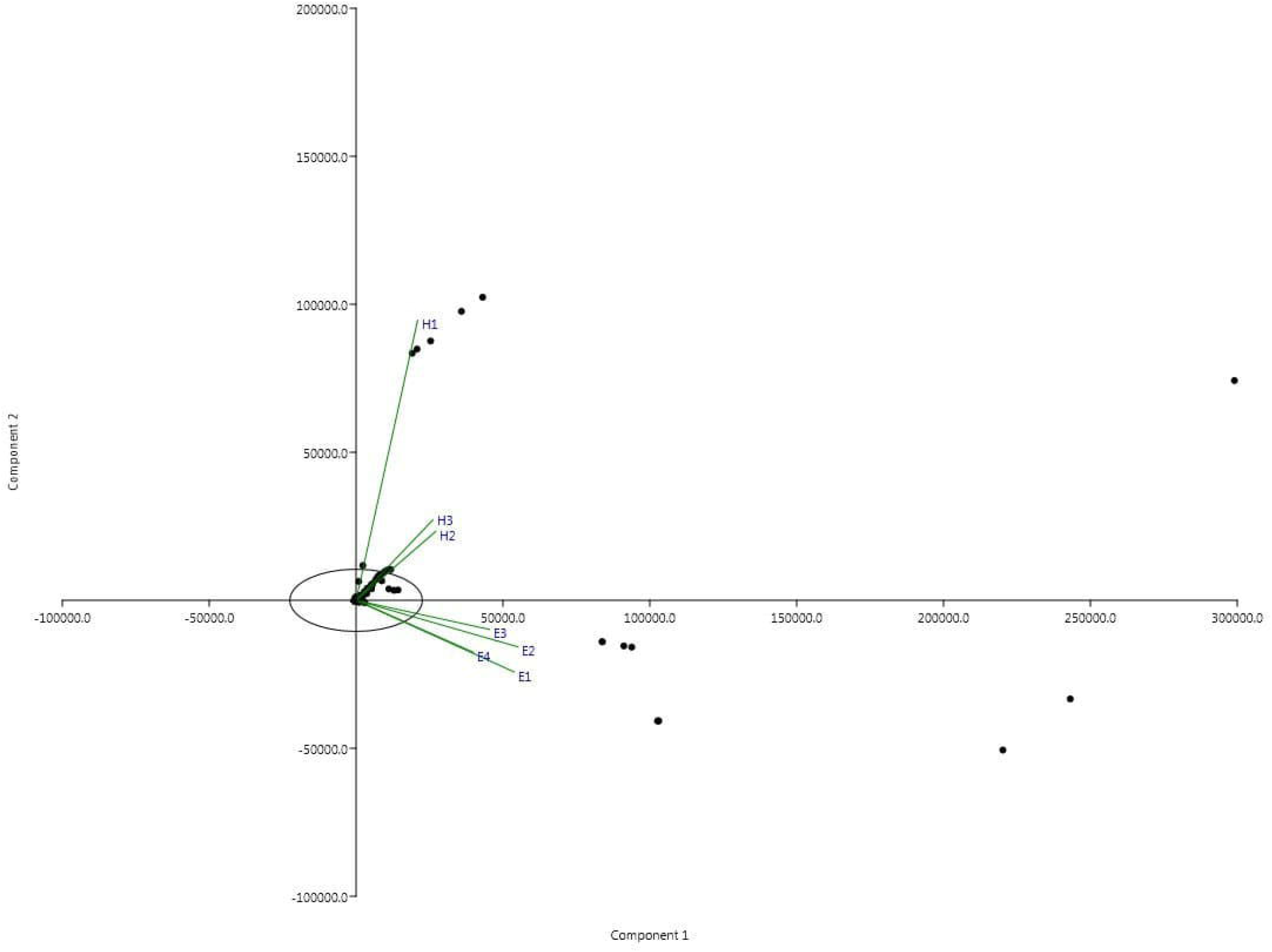
Principal component analysis (PCA) plot of the uterine proteome from healthy mares and mares with endometritis

**Figure 2 (a-g).**
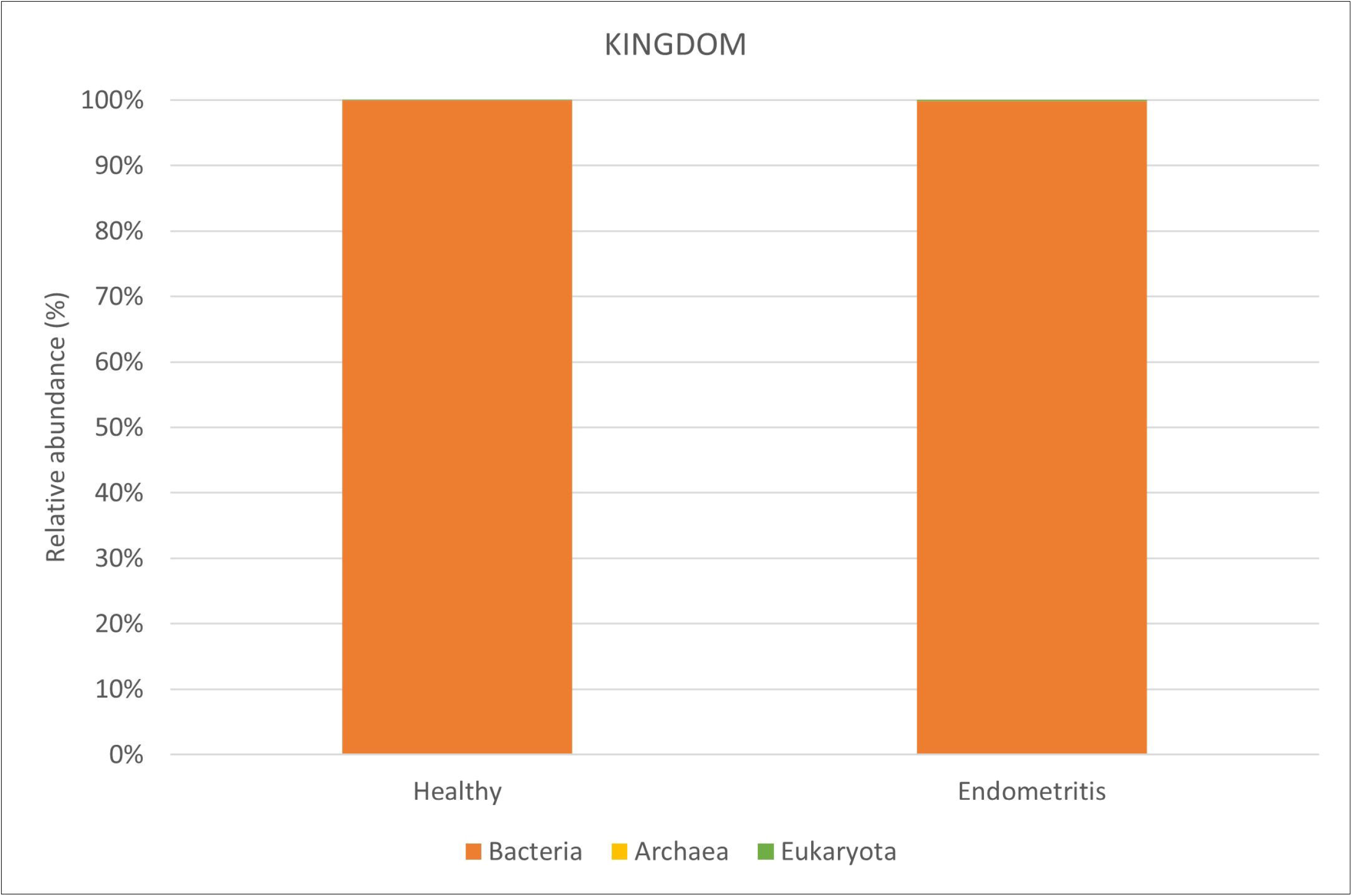
Taxonomical classification of the uterine proteome from healthy mares and mares with endometritis

At the genus level, *Brevibacillus* and *Paenibacillus* were more abundant in healthy mares, whereas *Escherichia, Salmonella*, and *Klebsiella* were more abundant in mares with endometritis (Figure 2f). In healthy mares, *Brevibacillus brevis* was the most abundant species, followed by *Brevibacillus choshinensis* and *Paenibacillus sp JDR-2*. However, in mares with endometritis, *Escherichia coli* was the most abundant species, followed by *Salmonella enterica* and *Klebsiella pneumoniae* (Figure 2g). The relative abundance of the various taxonomic levels in individual mares is shown in Supplementary Figure 1 (a-g).

## 4. Discussion

The detection of a diverse uterine microbiome in healthy mares in the present study, together with the findings of previous studies (Heil et al., 2018; Holyoak et al., 2022), strongly refutes the dogma of a sterile uterus in healthy mares. The composition of the uterine microbiome in the healthy mares in this study was mostly similar to that reported in a previous study in horses (Holyoak et al., 2022) and studies in other domestic animal species such as cows (Clemmons et al., 2017; Moore et al., 2017) and dogs (Lyman et al., 2019). Similar to the findings of Holyoak et al. (2022), proteobacteria and firmicutes constituted a vast majority of the uterine microbiome. However, there were some differences at the genus and species levels, which could be attributed to different geographical locations. Taken together, the results of these studies underscore the importance of metagenomic approaches in unravelling the diverse resident microbiomes in healthy organ systems that could not be detected using traditional microbial culture methods.

To our knowledge, the present study evaluated for the first-time differences in the uterine microbiome between healthy mares and mares with endometritis. The results showed notable differences in the uterine microbiome at various taxonomic levels between the two groups of mares. Interestingly, the major components of the uterine microbiome detected in healthy mares have been shown to be previously associated with defense mechanisms against pathogenic bacteria and fungi. Several studies have reported that *Brevibacillus* and *Paenibacillus* species produce a variety of peptides such as gramicidin and tostadin with documented broad spectrum antimicrobial activity (Wu et al., 2011; Song et al., 2012; Jianmei et al., 2015; Yang and Yousef, 2018). These findings suggest that the resident microbiome might have an important role in the uterine defense mechanism of mares and, consequently, their potential fertility.

Endometritis continues to be one of the major causes of infertility in mares and the findings of this study could potentially serve as a basis for novel diagnostic and therapeutic approaches to address this important problem. As suggested recently by Morrell and Rocha (2022), the interactions among various microbes rather than the presence of a single microbial species may play an important role in maintaining uterine health. The diverse uterine microbiome detected in the present study makes a strong case for potential inclusion of metagenomic profiling in the diagnostic work-up of infertile mares. The findings also have potential implications for treating mares diagnosed with endometritis. Traditionally, mares with endometritis are treated with intrauterine antimicrobials in conjunction with approaches to improve uterine clearance such as uterine lavage and systemic oxytocin treatment (LeBlanc, 2010; Morris et al., 2020). Based on the findings of this study, re-establishment of the resident microbiome using probiotics could be suggested as a potential adjunct to the traditional therapeutic approaches used for equine endometritis.

In summary, metagenomic analysis of uterine fluid samples helped unravel novel differences in the uterine microbiome between healthy mares and mares with endometritis. The findings of this study can serve as a basis for future research aimed at developing new diagnostic, preventative, and therapeutic approaches for equine endometritis.

---

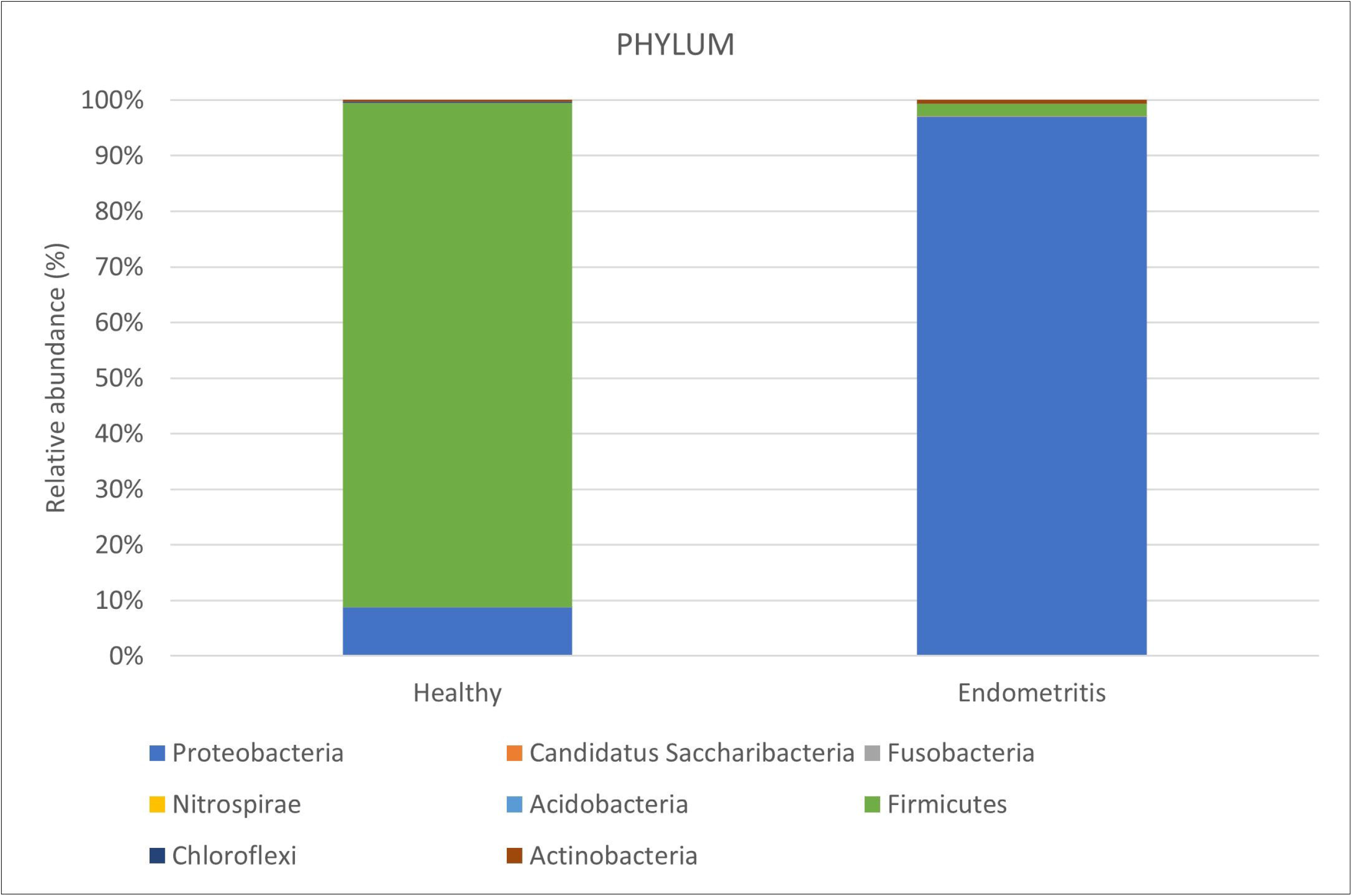

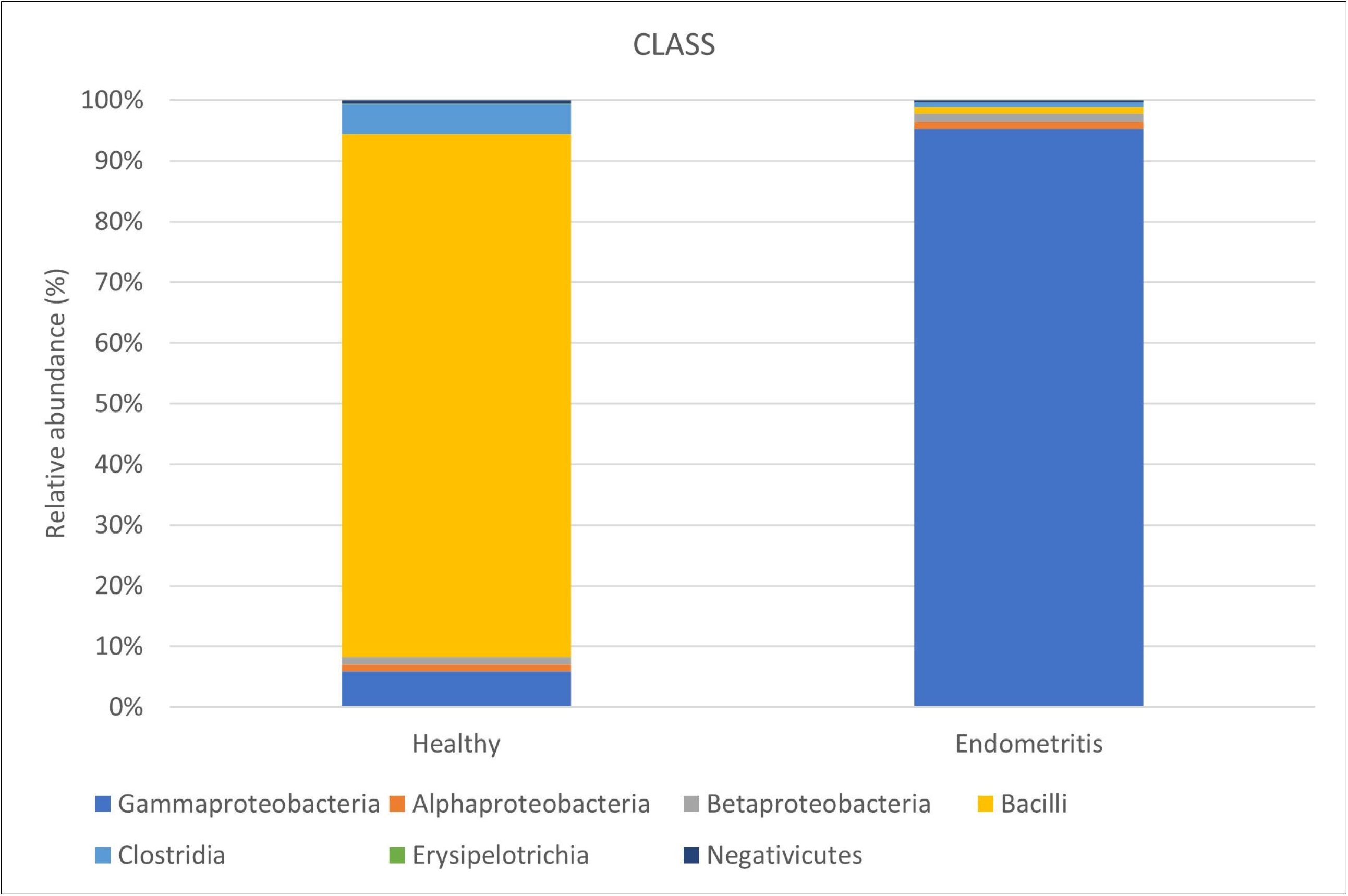

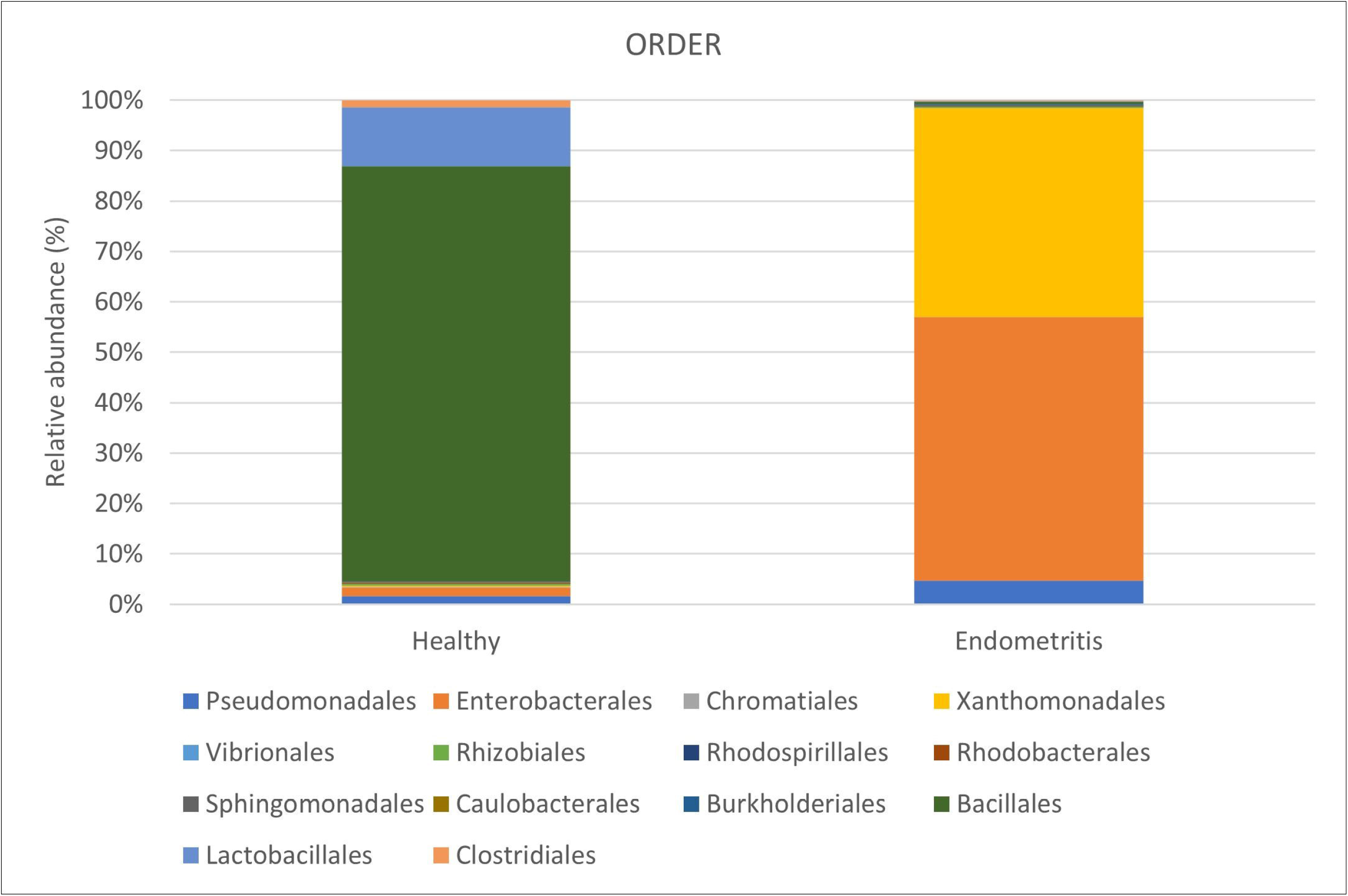

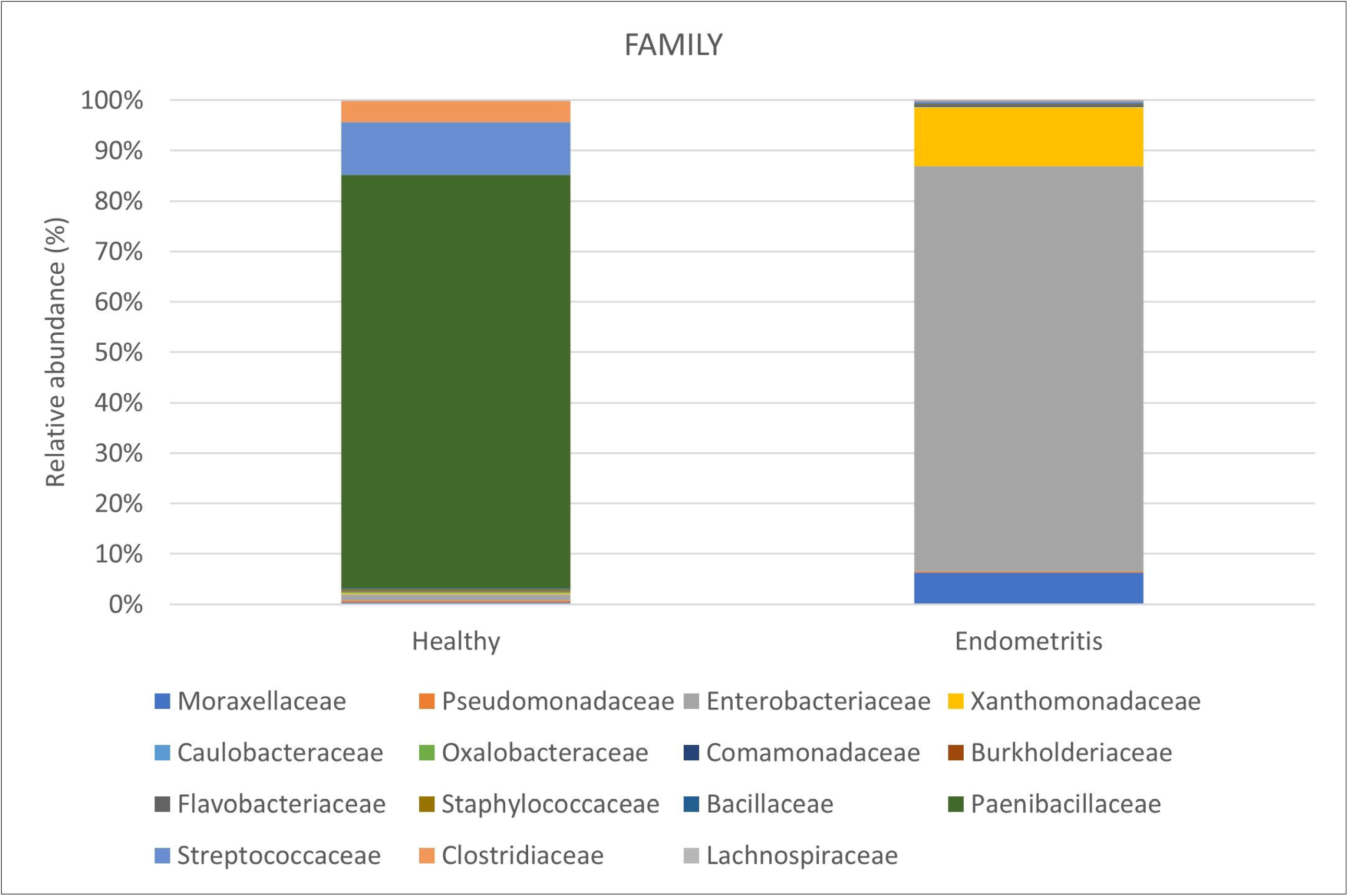

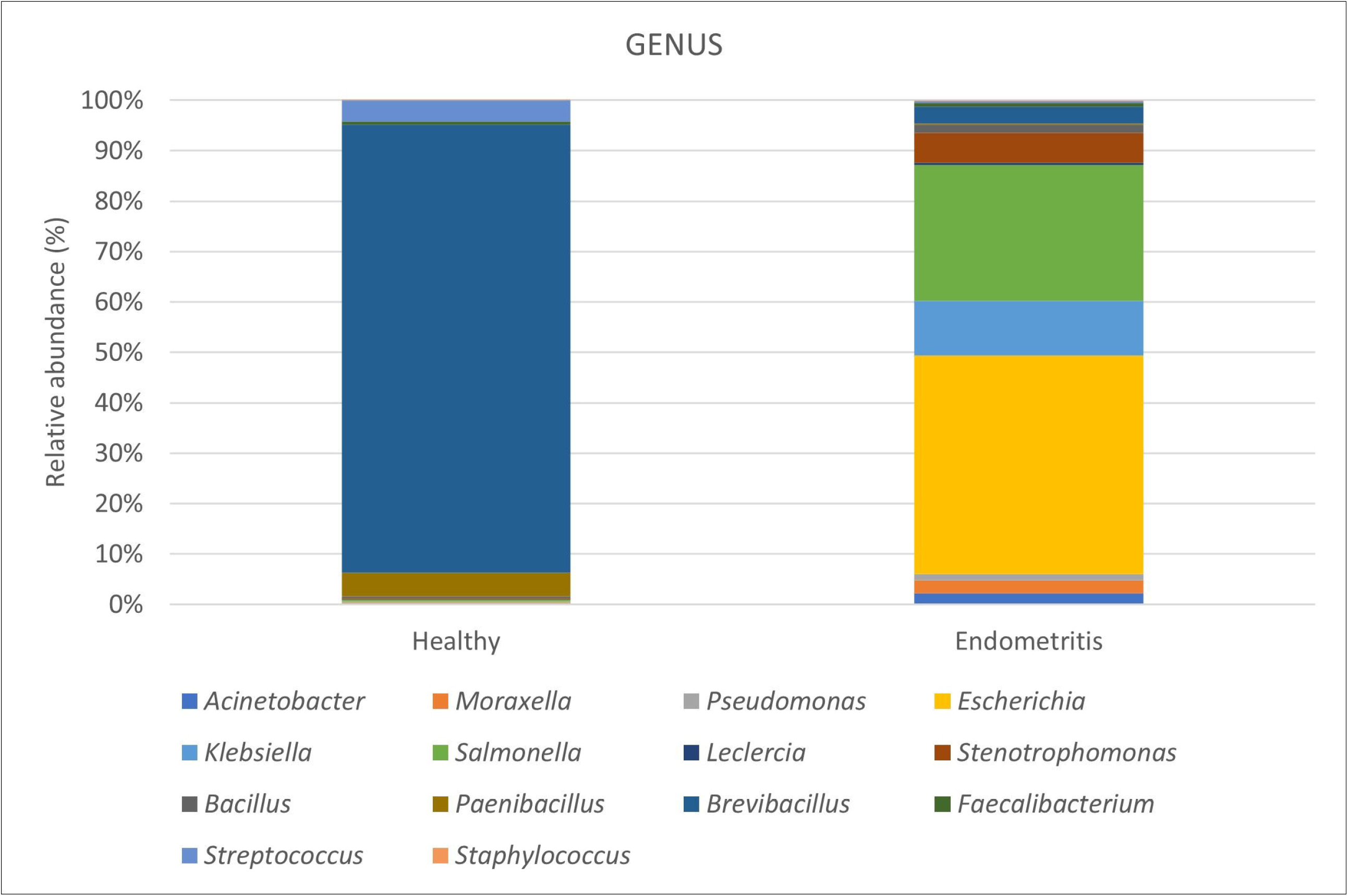

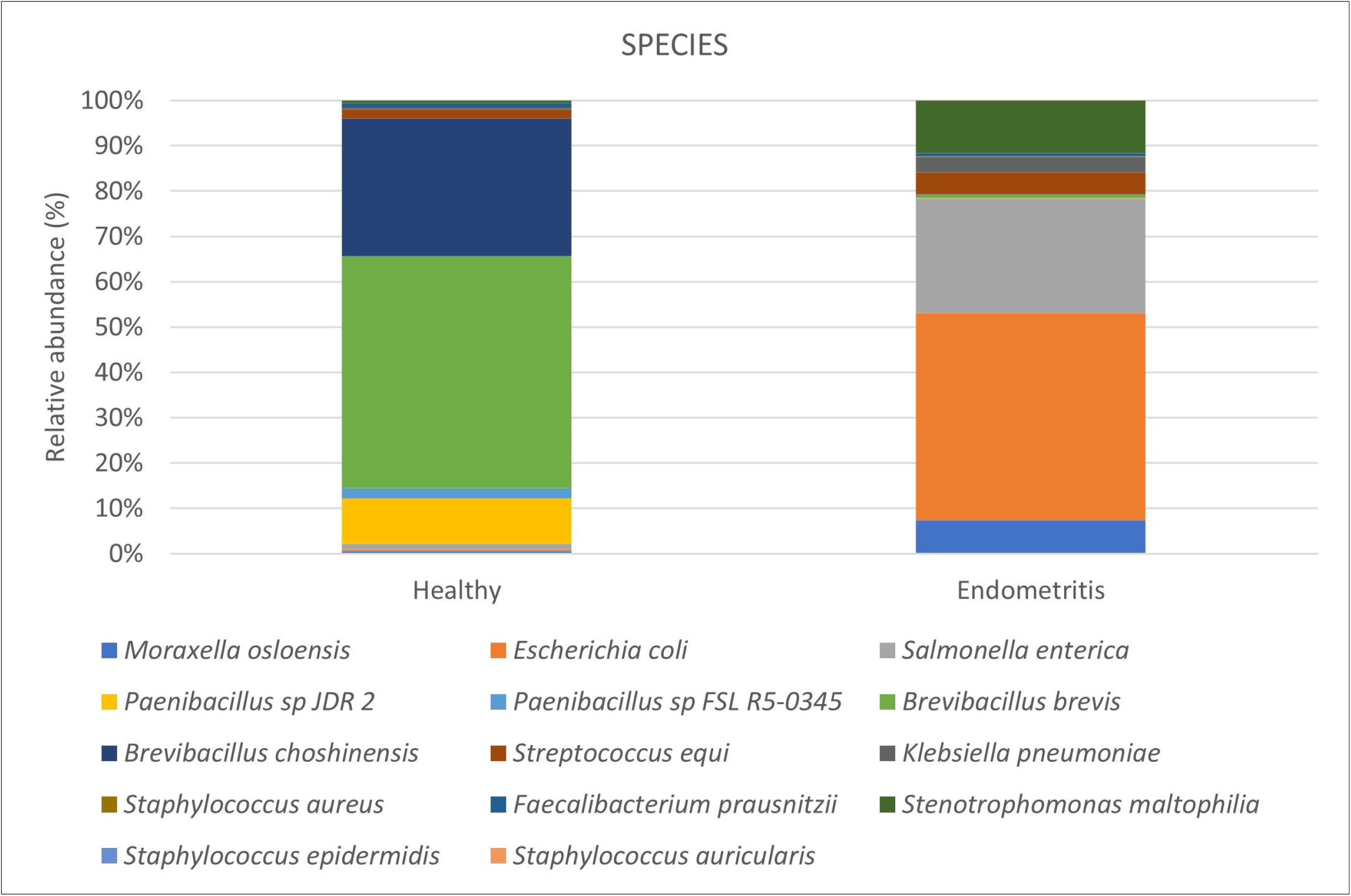

