## Supplementary figures and images for "Metagenomic analysis unravels novel taxonomic differences in the uterine microbiome between healthy mares and mares with endometritis"

### Supplementary Figure 1a.jpg

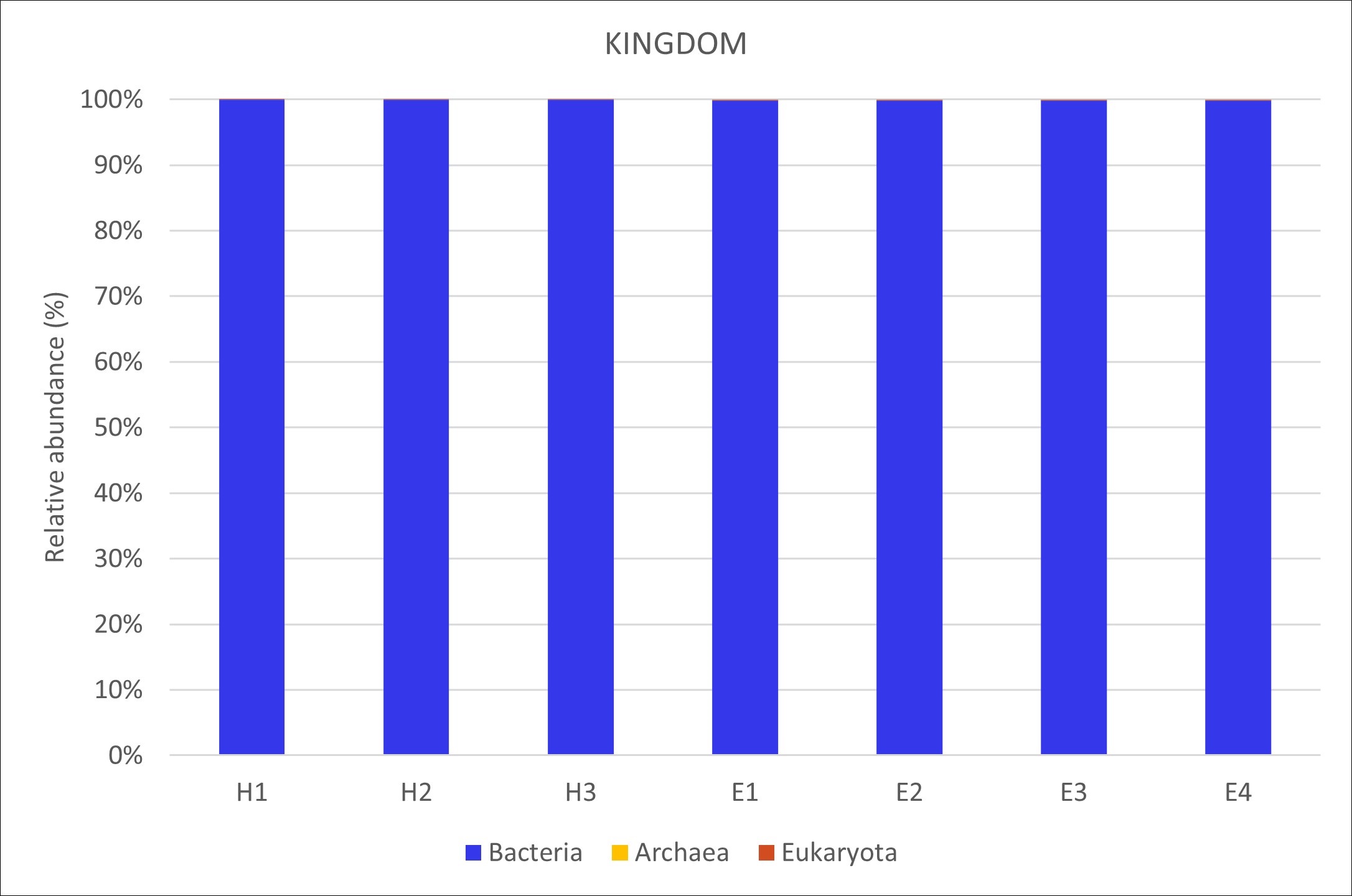

### Supplementary Figure 1b.jpg

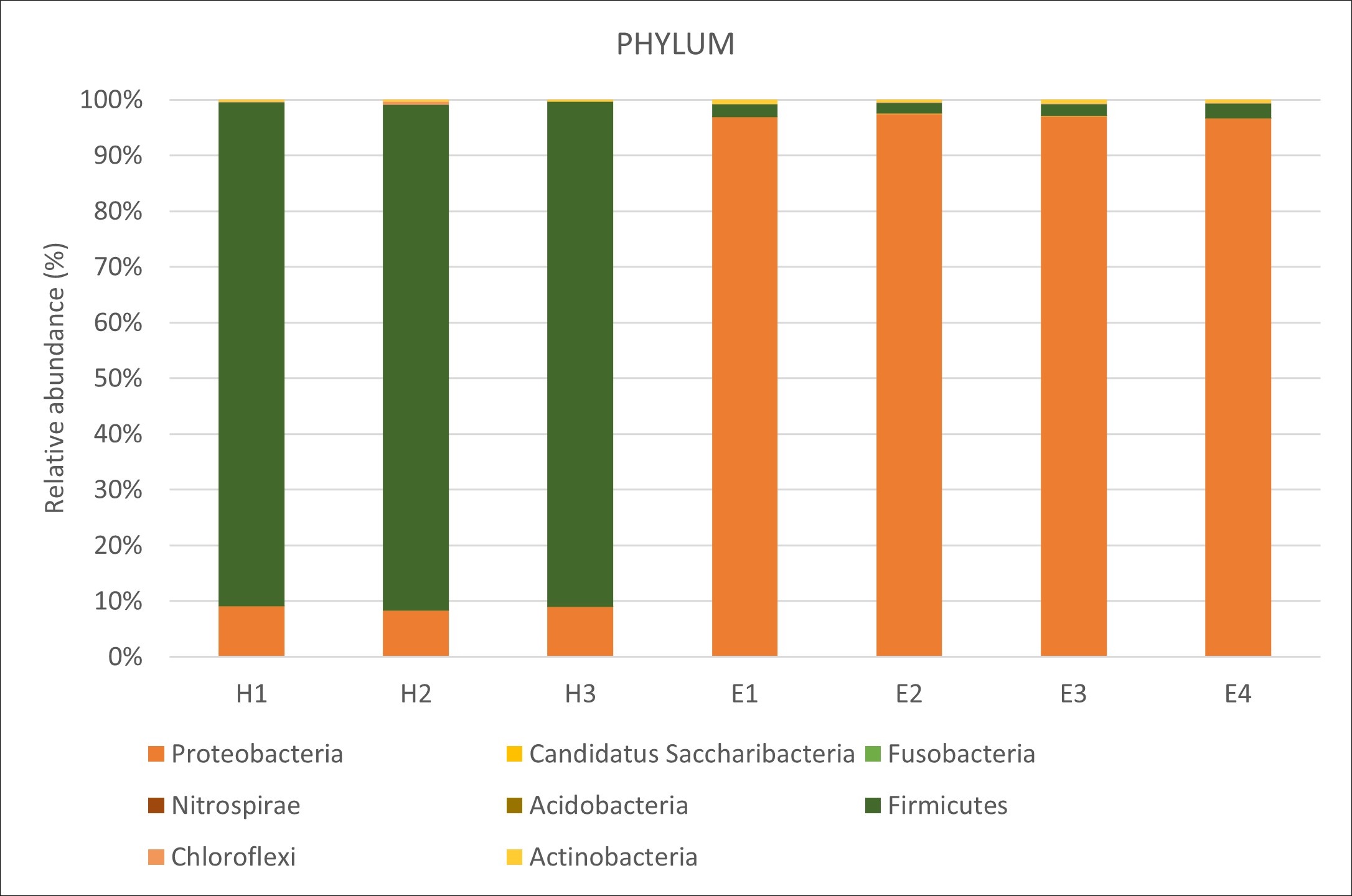

### Supplementary Figure 1c.jpg

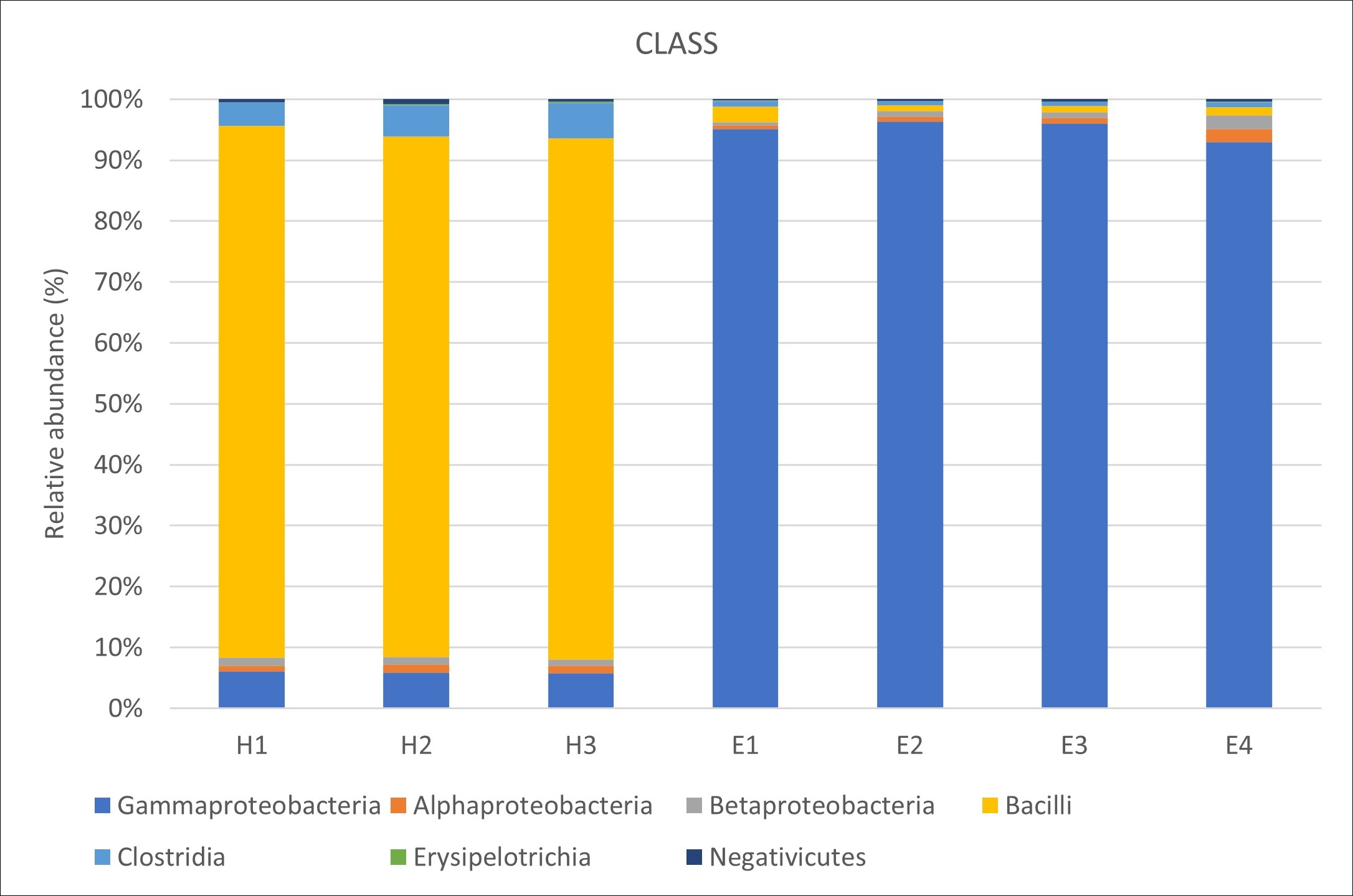

### Supplementary Figure 1d.jpg

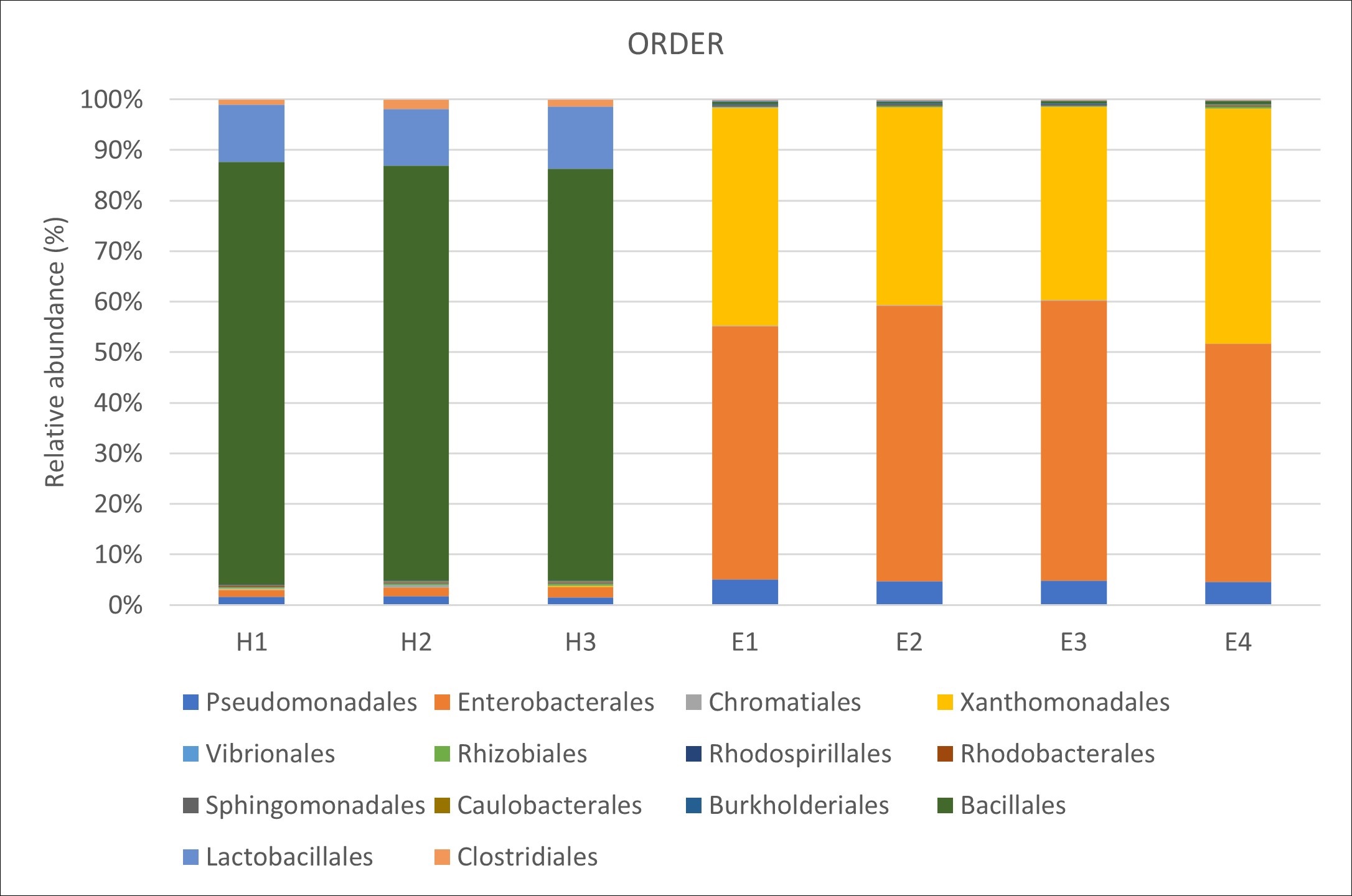

### Supplementary Figure 1e.jpg

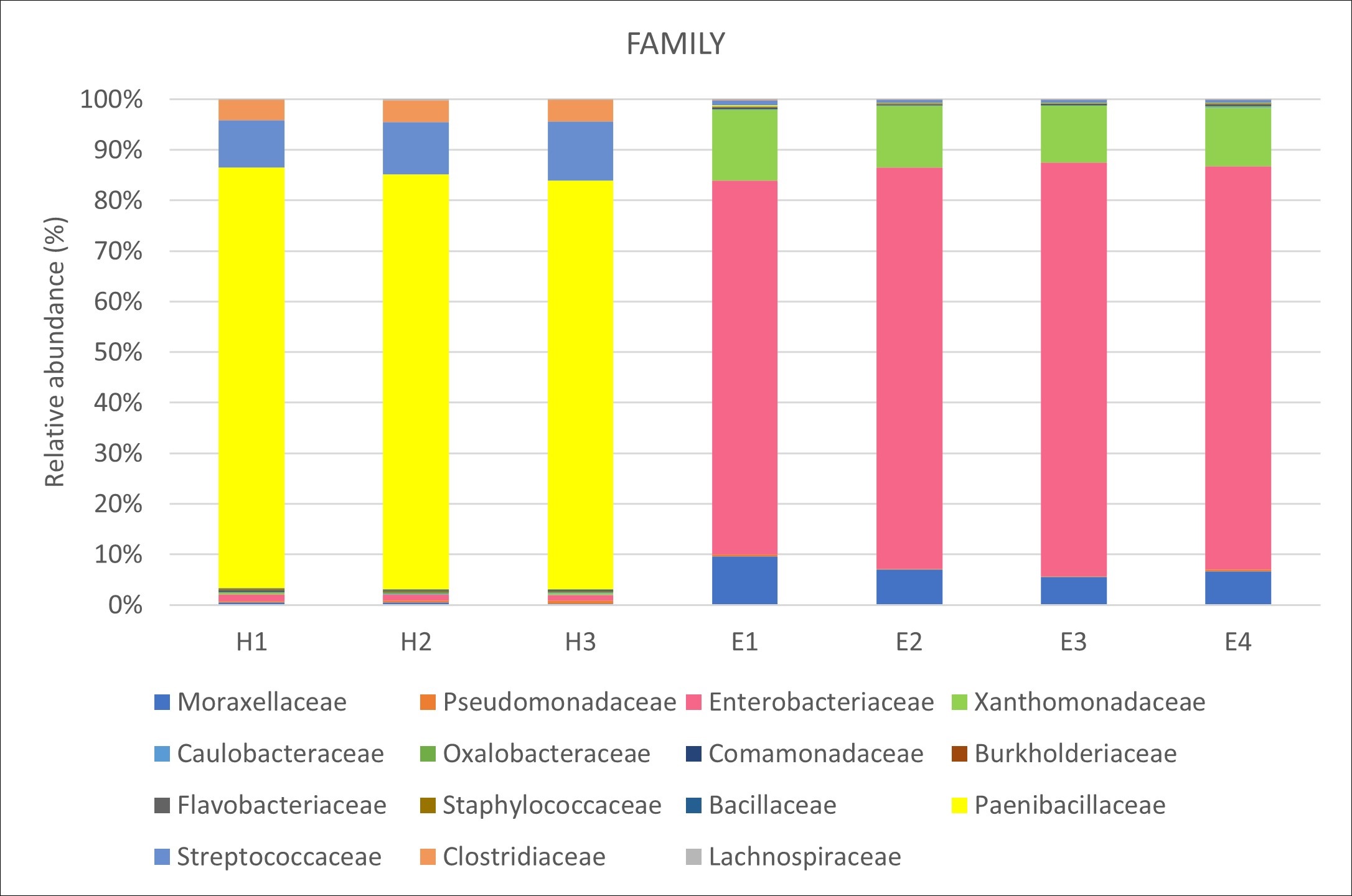

### Supplementary Figure 1f.jpg

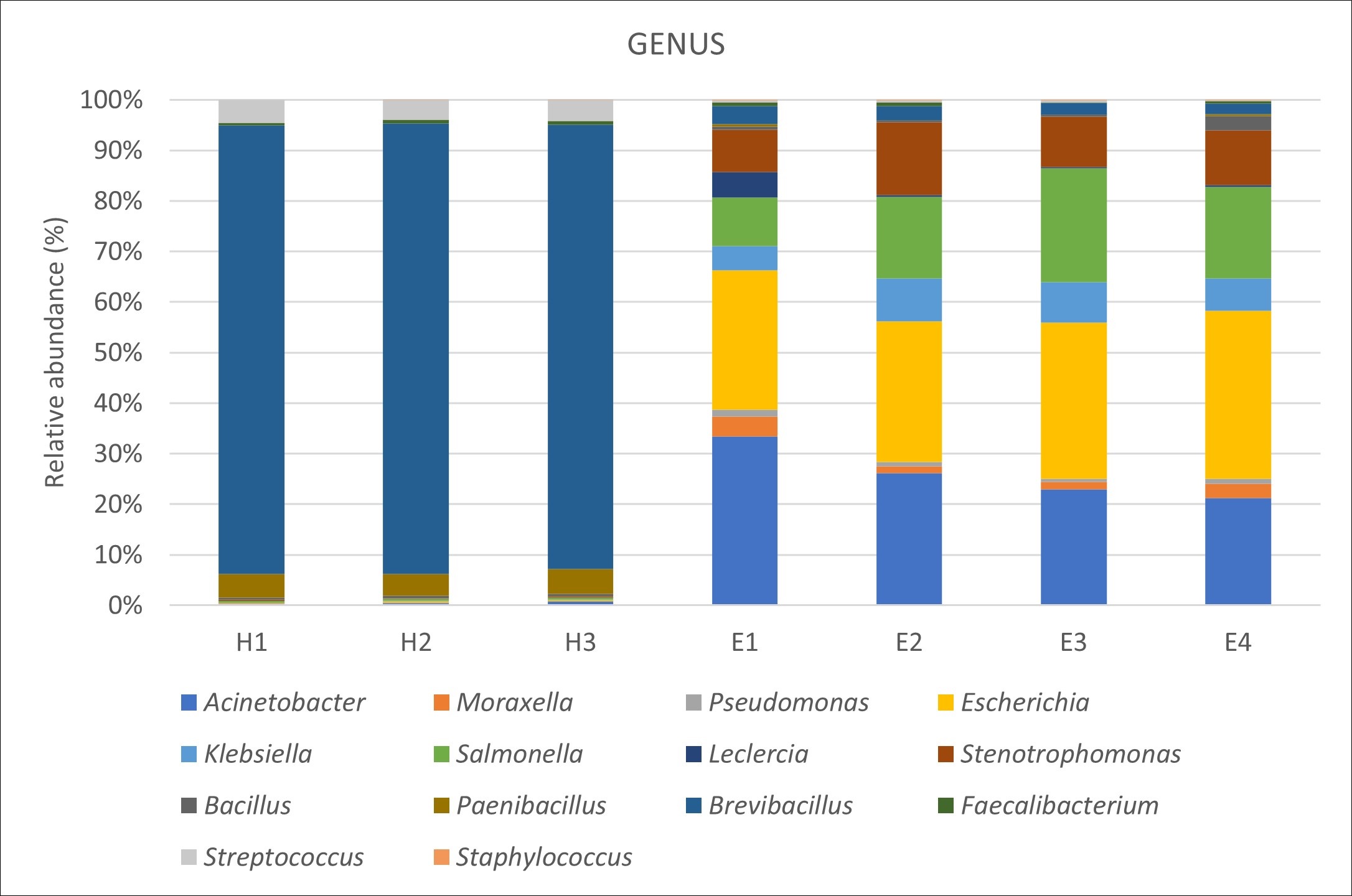

### Supplementary Figure 1g.jpg

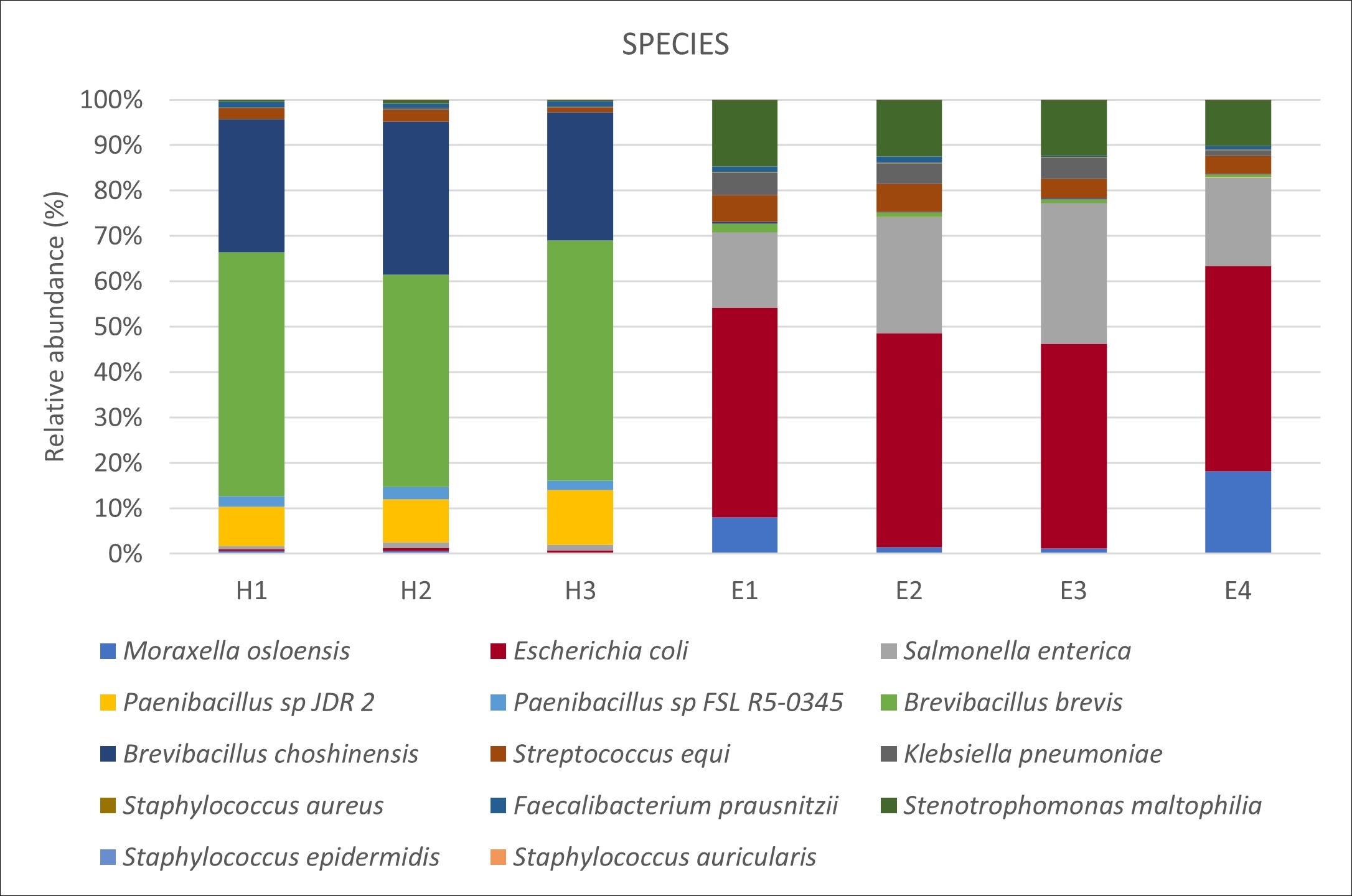
